# Removing head ganglia in amphibious centipedes unveils descending contribution to versatile locomotor repertoire

**DOI:** 10.64898/2026.04.02.716080

**Authors:** Kotaro Yasui, Emily M. Standen, Takeshi Kano, Hitoshi Aonuma, Akio Ishiguro

## Abstract

Understanding how animals produce a versatile locomotor repertoire requires unraveling the interplay between higher centers, decentralized locomotor circuits, and sensory feedback. However, the principles governing their integration remain elusive. We investigated amphibious centipedes through stepwise neural lesions and neuromechanical modeling. Behavioral experiments revealed that while decentralized circuits autonomously generate coordination, the brain and subesophageal ganglion provide situational flexibility, such as modulating trunk undulation and initiating leg folding. Integrating these findings, our model demonstrated how higher centers selectively inhibit or release lower circuit dynamics. Simulations verified that varying only a few descending control parameters reproduces transitions between slow walking, fast walking, and swimming. This work may capture the essence of the locomotor circuitry that harnesses decentralized self-organization to coordinate the body’s large degrees of freedom.

---

Animals exhibit adaptive locomotor behaviors in response to their environment, which is crucial for their survival. For instance, they realize higher locomotion speed, mostly associated with gait changes (*1, 2*), when they escape from predators, and different locomotor patterns such as walking and swimming are selectively employed when they encounter different environmental substrates (*3–5*). Such situation-dependent locomotion, commonly observed across animal species, is achieved by appropriately selecting which body parts to move for propulsion and flexibly coordinating motor output patterns between them. Observing animals across a variety of environments, therefore, allows us to decode locomotor circuits capable of shaping adaptive coordination and to understand the control principles underlying behavioral flexibility in animals.

Neurobiological studies, using various model animals, have long suggested that locomotor movements are generated through the interplay between the higher nervous system (i.e., the brain) and the lower nervous system (i.e., spinal cord in vertebrates, ventral nerve cord in invertebrates). The higher center is responsible for selecting locomotor modes (*6–8*), such as walking, swimming, or flying, and for tuning the locomotion speed and direction of the movement (*9, 10*). In contrast, the lower nervous system plays a major role in producing the actual motor output patterns for locomotion. Specifically, distributed neural circuits along the lower nerve cord, referred to as central pattern generators (CPGs) (*11–15*), are capable of generating rhythmic neuronal activities for locomotor movements. While the CPGs can produce basic neuronal rhythms without receiving any rhythmic inputs from the higher centers and peripheral sensory organs, sensory feedback loops are also suggested as key mechanisms for adjusting the rhythmic patterns of CPGs in response to environmental changes and thus shaping the coordination between different body parts (*16–21*). Integrating the brain, lower nerve cord, and sensory feedback is important to illustrate the entire schematic of the control circuitry underlying flexible locomotor behavior in animals (*22*). While a few recent studies have employed computational modeling to explore the flexible locomotor control for behavioral versatility (e.g., in salamanders (*23*) and insects (*24*)), how the higher centers and the lower locomotor circuits are functionally integrated in animals largely remains elusive.

Here we focus on an amphibious centipede (*Scolopendra subspinipes mutilans*). The centipede exhibits versatile body and limb coordination for locomotion (*25, 26*): On land, it walks slowly by propagating a wave of leg movement (Fig. 1A), whereas, in water, it swims with a traveling wave of trunk undulation (Fig. 1B). During fast walking, it combines a swimming-like wave of trunk undulation with that of leg movement (Fig. 1C). Anatomical studies suggest that this locomotor repertoire is generated via the central nervous system which is composed of head ganglia and a series of segmental ganglia (Fig. 1D) (*27, 28*). As it was reported that centipedes can walk without a head (*29*), the lower locomotor circuits (segmental ganglia) are likely to be resilient in producing the walking behavior without signals from the brain. Accordingly, taking advantage of the relatively simple neural structure (i.e., serially connected ganglia) compared to vertebrates and the capacity of the system to endure neuronal lesions, amphibious centipedes can be a suitable model to investigate how the higher centers interact with the lower locomotor circuits to realize a versatile locomotor repertoire in response to different environmental situations.

**Figure 1:**
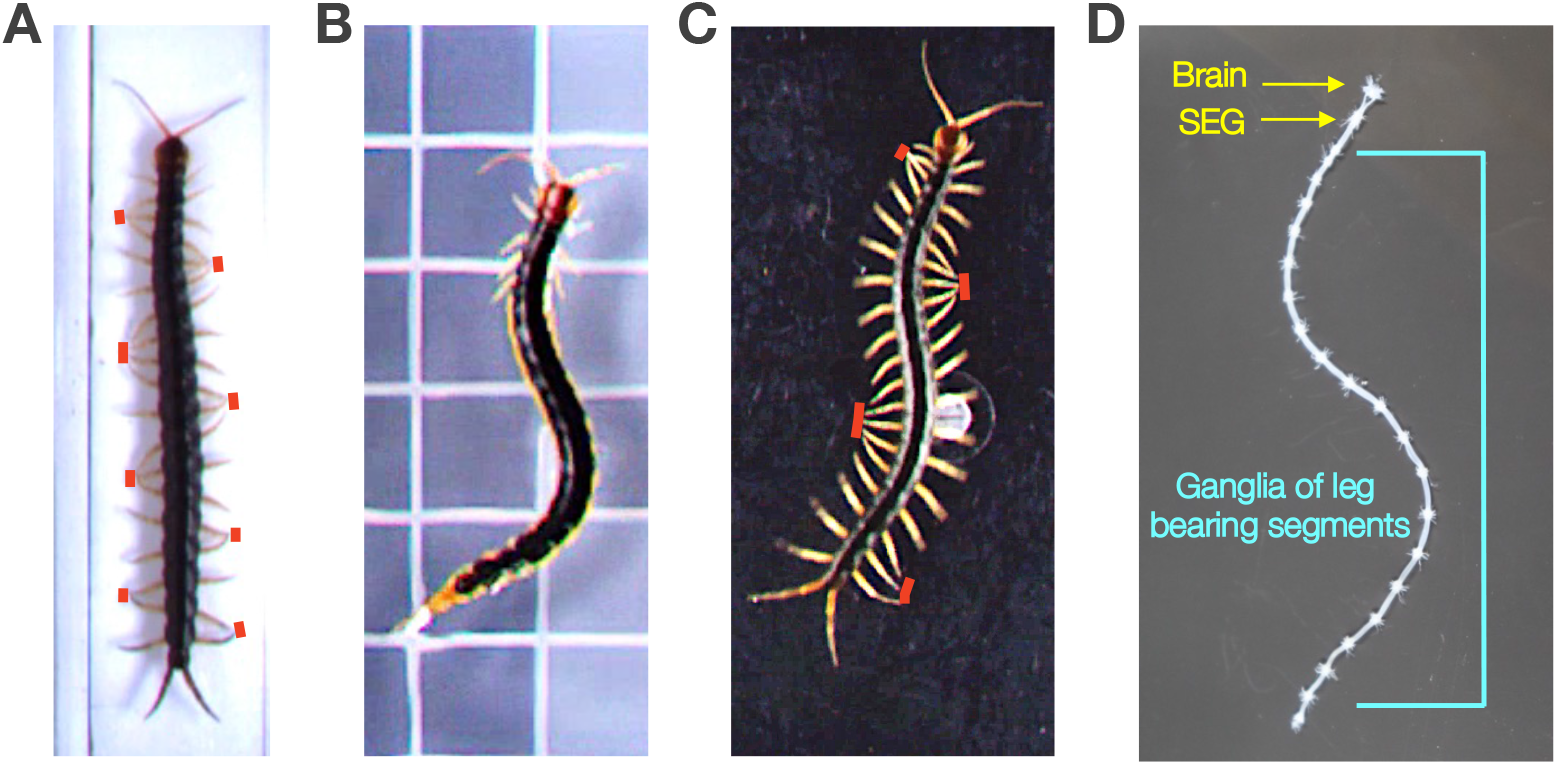
Locomotor patterns and central nervous system of the centipede, *Scolopendra subspinipes mutilans*. (A) Walking at slow speed. (B) Swimming in water. (C) Walking at high speed. (D) The brain, subesophageal ganglion (SEG), and ganglia of leg-bearing segments. Red bars in (A) and (C) indicate the legs in contact with the ground.

Thus far, we have adopted a synthetic approach (*30–32*) for understanding the essence of locomotor circuits in centipede locomotion, using mathematical models based on behavioral findings. Our previous models proposed two hypotheses regarding the lower locomotor circuits responsible for walking that require further testing in the biological system:

1. The ‘*ground contact*’ hypothesis suggests that the inter-limb coordination pattern for walking can be generated through a decentralized control system using mechano-sensory feedback based on the ground contact of the legs (*33–35*).
2. The ‘*trunk-limb*’ hypothesis where the coordinated pattern between the trunk and legs during fast walking can be established through bidirectional local sensory feedback between the trunk and limbs (*36*).

Although our previous models successfully replicated slow and fast walking patterns in simulations that did not require a brain, it has not been confirmed in living animals if the resultant gaits can be produced with only lower locomotor circuits (without higher brain centers). By removing brain sections from living centipedes, this paper aims to verify the ‘*ground contact*’ and ‘*trunk– limb*’ hypotheses (*33–36*) and in doing so identify the functional significance of different brain components. Brain is important for the walk–swim transition. Our previous work transecting centipedes mid-body has shown that initiating swimming requires a descending command from the higher centers and that local sensory feedback from the legs may override this swimming command to elicit walking at the local segment level (*25*). However, this previous study used centipedes whose ventral nerve cord was transected at the middle of the body, allowing the descending signal from the brain to occur in the anterior sections of the animal. Without isolating independent brain regions, the role of the fore brain ganglia and the subesophageal ganglia (SEG), in controlling different locomotor patterns and how the descending signal interacts with lower locomotor circuits remains unclear.

In this study, we performed stepwise removal of the brain and SEG of the centipede to identify the differing contributions to locomotor control of the brain regions. The experimental results reported below support the ‘*ground contact*’ and ‘*trunk-limb*’ hypotheses proposed in our earlier paper and suggest that leg folding during swimming and trunk undulation during fast walking and swimming may be controlled by the descending signals from the higher centers via hierarchical interactions between them. Combining these new experimental data with the essence of previous models, we propose a single, comprehensive mathematical model of centipede locomotion. Our proposed model is the first to reproduce versatile locomotor behaviors (including the transition between slow walking, fast walking, and swimming) through simple descending control from the higher centers interacting with lower locomotor circuits. Our results provide a foundation for understanding the control principles underlying situation-dependent locomotion in animals.

## Results

### Behavioral experiments

To understand how the higher centers interact with the lower locomotor circuits in situation-dependent locomotion, we performed behavioral experiments in terrestrial and aquatic environments, using centipedes whose brain and then SEG were surgically removed (for details, see Materials and Methods). The higher centers in centipedes consist of the brain and the SEG (Fig. 1D) (*27, 28*) and may contribute to the control of locomotion as is seen in some less elongate arthropods (*37–39*).

First, we surgically cut the connectives between the brain and SEG in the centipedes (Fig. 2A, Supplementary Movie S1) and observed their behavior. On land, the brainless centipede exhibited coordinated walking with a straight body and a traveling wave of leg movement, similar to the slow walking pattern of intact centipedes (Fig. 2B). In water, the locomotor outcome was more varied (see Table 1). The four possible combinations of leg folding and body oscillation appeared in the brainless animals. Interestingly, in roughly half of the trials, rhythmic trunk undulation (46%), and leg folding (57%) appeared; however, these two behaviors were not always coupled (Fig. 2C). These results suggest that trunk undulation and leg posture are decoupled and that the propensity for rhythmic trunk undulation decreases in both terrestrial and aquatic environments when the brain is lost, however, with the SEG intact brainless centipedes still retain the ability to initiate leg folding and swimming-like trunk undulation.

**Table 1:**
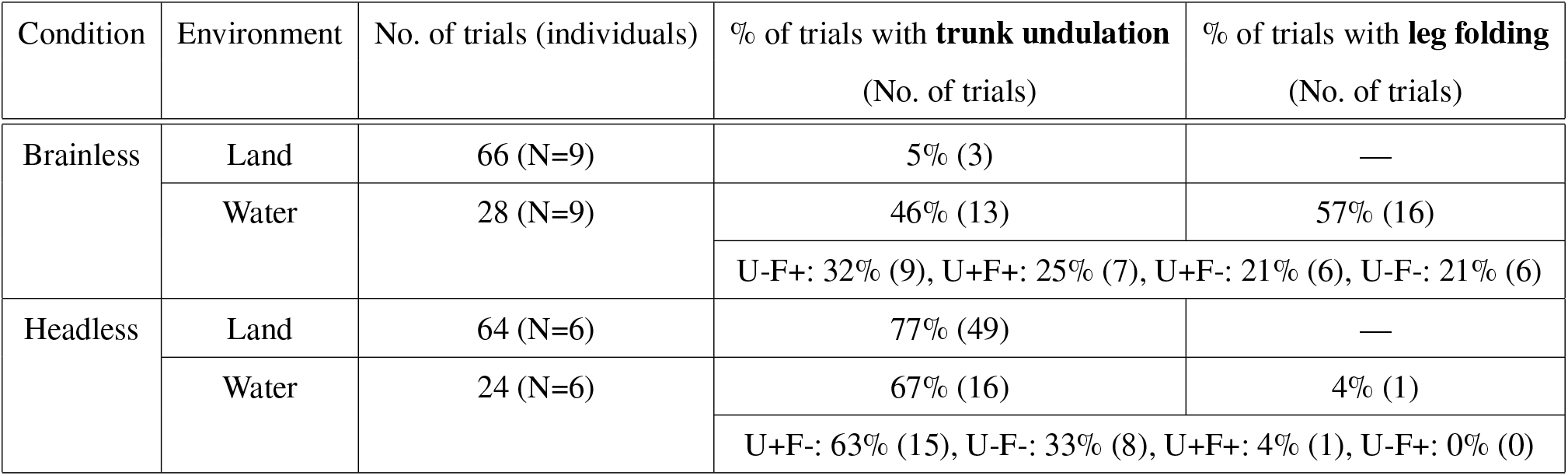
Summary of observed behaviors of centipedes on land and in water following surgical removal of the brain and head. For results in water, behavioral combinations are shown in the bottom row. ‘U’ and ‘F’ denote trunk undulation and leg folding, respectively, and ‘+’ and ‘–’ indicate the presence and absence of each behavior.

**Figure 2:**
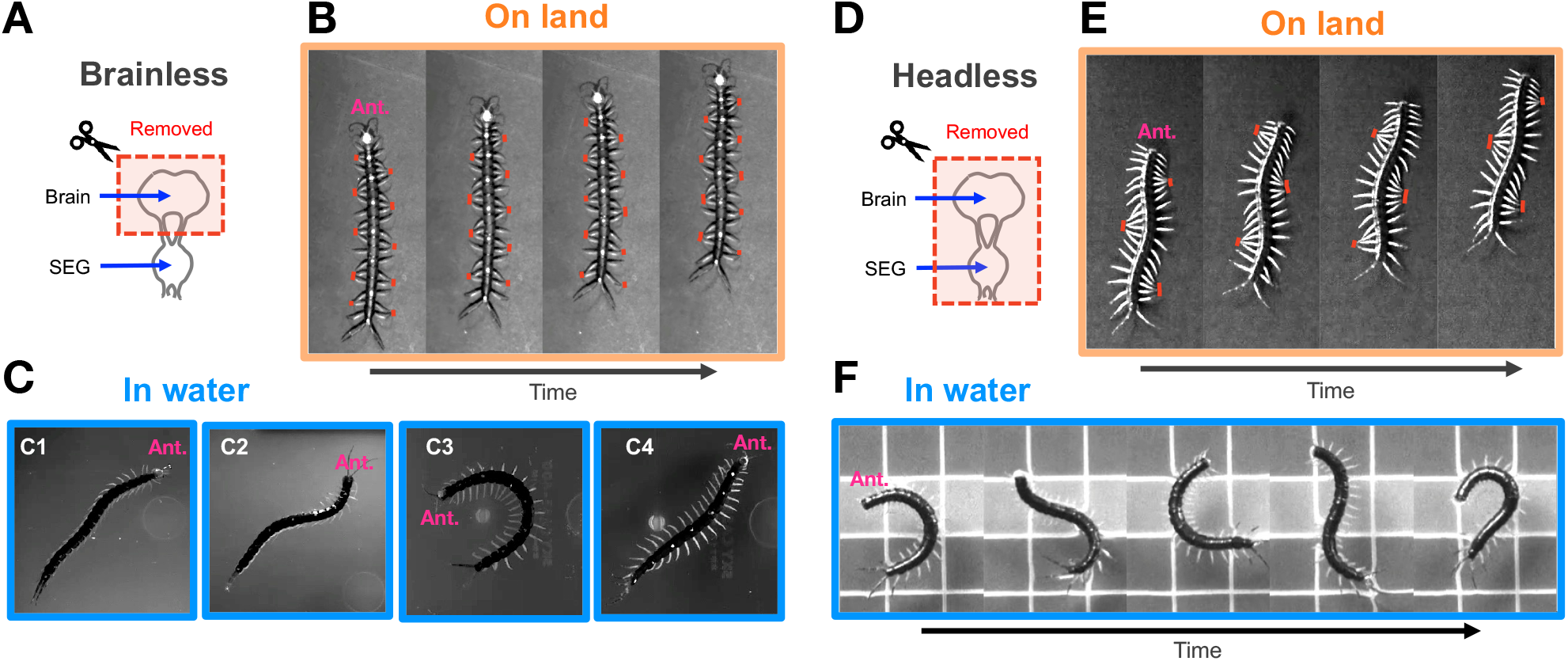
Behavioral observations after surgically removing the head ganglia from centipedes. (A to C) Behaviors after the brain was removed (the head is still present). (D to F) Behaviors of headless centipedes (i.e., removal of the brain and SEG). (A) and (D) illustrate the surgically removed part of the head ganglia. Red bars in (B) and (C) indicate the legs in contact with the ground. The abbreviation ‘Ant.’ denotes the anterior direction.

Next, we removed both the brain and SEG by removing the head from the centipedes and observed their behavior (Fig. 2D, Supplementary Movie S2). It should be noted that we used different individuals for this experiment to minimize the effect of physiological fatigue and damage after the surgery of brain removal. On land, the headless centipede primarily exhibited the fast walking pattern seen in intact centipedes; this consisted of a traveling wave of leg movement and trunk undulation (77% of trials Fig. 2E). Indeed, headless centipedes exhibited significantly faster walking speeds (1.00±0.39 body lengths per second [BL/s], N=6) compared to brainless ones (0.37±0.22 BL/s, N=9; P=0.0028, Mann-Whitney U test). Data are presented as mean ± SD of per-individual averages. On occasion, but rarely, headless centipedes could exhibit a slow walking pattern with little to no trunk undulation. Usually, the slow walk transitioned into a fast walking pattern quite quickly. In water, the headless centipedes held their legs in an extended position and often exhibited rhythmic trunk undulation (Fig. 2F, Table 1). Note that they did not immediately initiate the swimming-like trunk undulation upon entering the water; rather, they started it after some time had passed, and mostly the magnitude of trunk undulation increased gradually (see Supplementary movie). These results suggest that the rhythmic trunk undulation is inhibited by the activity of an intact SEG both in terrestrial and aquatic environments. The data also suggest that sensory feedback magnitude may build with increasing motion and set the magnitude of the undulation. Because leg folding behavior for swimming almost completely disappears after the loss of SEG, this data also suggests that SEG shares control of leg position with the brain.

Considering the above-mentioned behavioral findings, we hypothesize that the following control mechanisms underpin centipede locomotion (Fig. 3):

**Figure 3:**
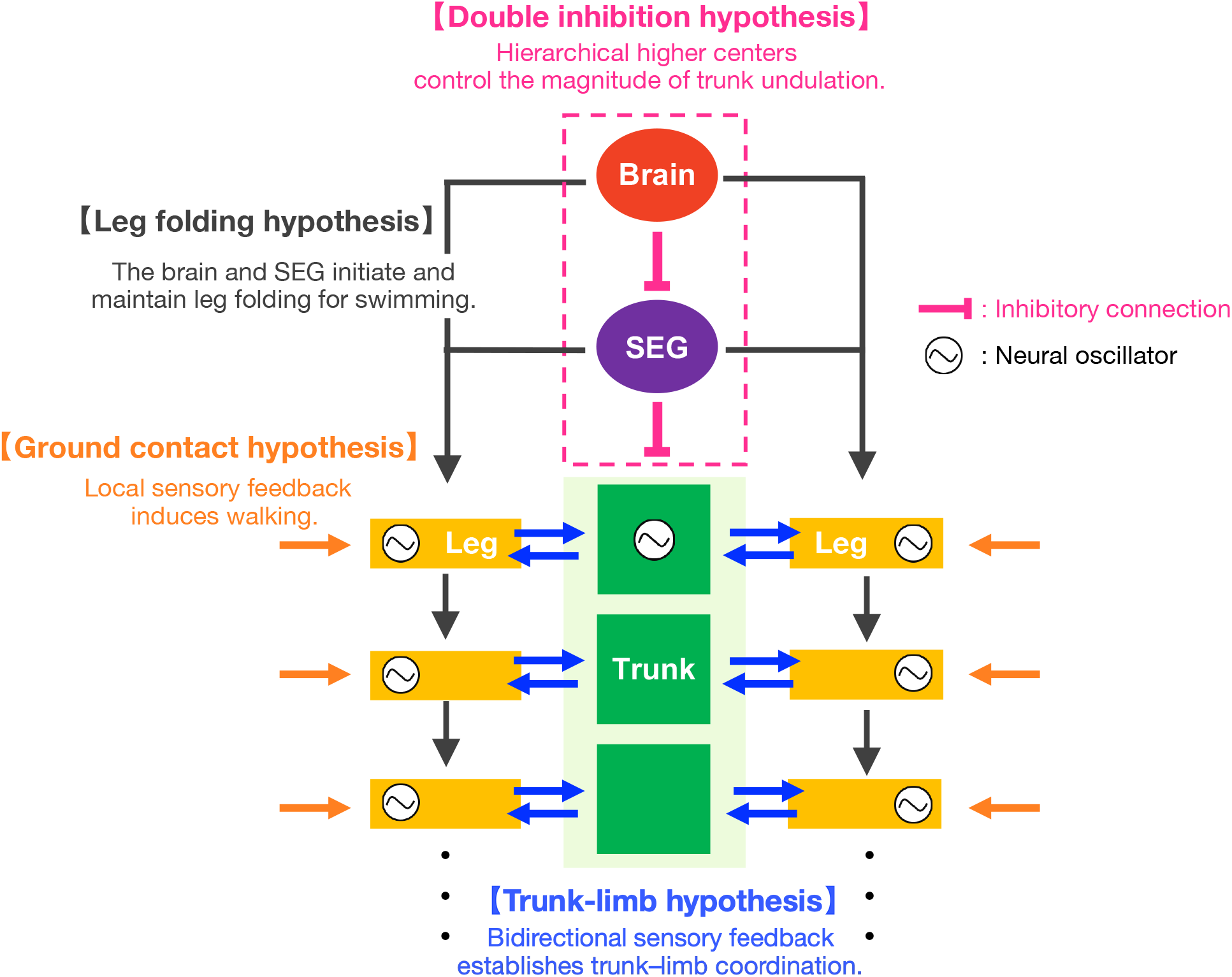
Schematic of the hypothesized hierarchical control circuits in centipede locomotion. This integrated model combines decentralized locomotor circuits proposed in our previous studies with descending control mechanisms newly proposed in this work. The ‘ground contact’ (orange) and ‘trunk–limb’ (blue) hypotheses represent the decentralized control via local sensory feedback that was proposed in our earlier models to explain inter-limb and trunk-limb coordination for walking. Building on this, the current study introduces the ‘double inhibition’ hypothesis (pink) to describe how the brain releases the inhibition on trunk undulation from subesophageal ganglion (SEG), alongside the ‘leg folding’ hypothesis (black) which posits that additive descending signals from both the brain and SEG initiate the leg posture during swimming.

(3) The ‘*double inhibition*’ hypothesis, where the SEG inhibits the rhythmic trunk undulation, and the brain initiates and controls the magnitude of trunk undulation during fast walking and swimming by inhibiting the activity of the SEG.
(4) The ‘*leg folding*’ hypothesis, where the brain and SEG signals initiate and maintain leg folding for swimming and these signals are additive.

### Mathematical modeling

To test the hypothesized control mechanisms, we built a simple two-dimensional neuro-mechanical model of the centipede with 20 pairs of legs (Fig.4A). The centipede body is modeled using a mass-spring-damper system. To represent the leg-tip motion on a horizontal plane, each leg is assumed to be equipped with a rotational actuator at its base that swings the leg forward and backward, as well as a linear actuator that allows adjustment of the leg’s axial length. Mimicking the leg-tip trajectory of living centipedes, the linear actuator actively shortens the leg during the stance phase to achieve a straight-line motion of the leg tip parallel to the body axis, whereas the leg length remains constant during the swing phase (see Eq. 11 in Materials and Methods). To describe the neuronal rhythms of the leg (i.e., CPG) during walking, we implemented a phase oscillator for each leg. On land, the leg tip is assumed to be in the swing phase (lifted off the ground) when the leg oscillator phase 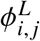 is 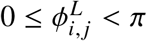, and in the stance phase (contacting the ground) when 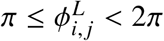. Between each body segment, a rotational actuator is assumed, enabling the generation of trunk bending motion. These rotational and linear actuators in the legs and trunk are controlled to achieve the target joint angles and leg lengths specified by the outputs of the locomotor circuits (see the following section for details).

#### Control circuits for trunk motion

To generate the flexible coordination between the body segments during locomotion, our model describes the target joint angle of the *i*th trunk segment 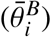 as follows:

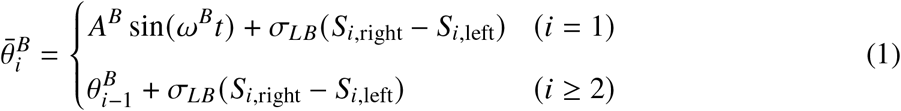

We assume that the trunk at the first body segment (*i* = 1) bends periodically with an amplitude *A*^*B*^ and a angular frequency of *ω*^*B*^ at time *t*, which corresponds to the intrinsic oscillator rhythm for undulatory movement. Then, the trunk bending initiated from the first segment propagates posteriorly by each segment (*i* ≥ 2) following the actual angle of its nearest anterior segment 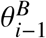 (Fig.4B). We employed this control scheme because it is suitable for minimally describing the nature of undulatory locomotion control (see (*40*) for details). In addition to this basic generator of undulation, we have proposed a local sensory feedback mechanism for trunk-limb coordination (‘***trunk–limb*’ hypothesis**) (*36*), described in the second terms of the right-hand side in Eq. 1 (Fig. 4C). Here, *σ*_*LB*_ is the positive constant which represents the strength of the sensory feedback, and *S*_*i*,left_ and *S*_*i*,right_ denote the ground contact sense of the left and right legs (see Eq. 15 in Materials and Methods). This sensory feedback was designed based on the kinematic relationship between the trunk and limbs during fast walking (Fig. 1C) and works such that the trunk bends into a concave shape toward the grounded leg.

**Figure 4:**
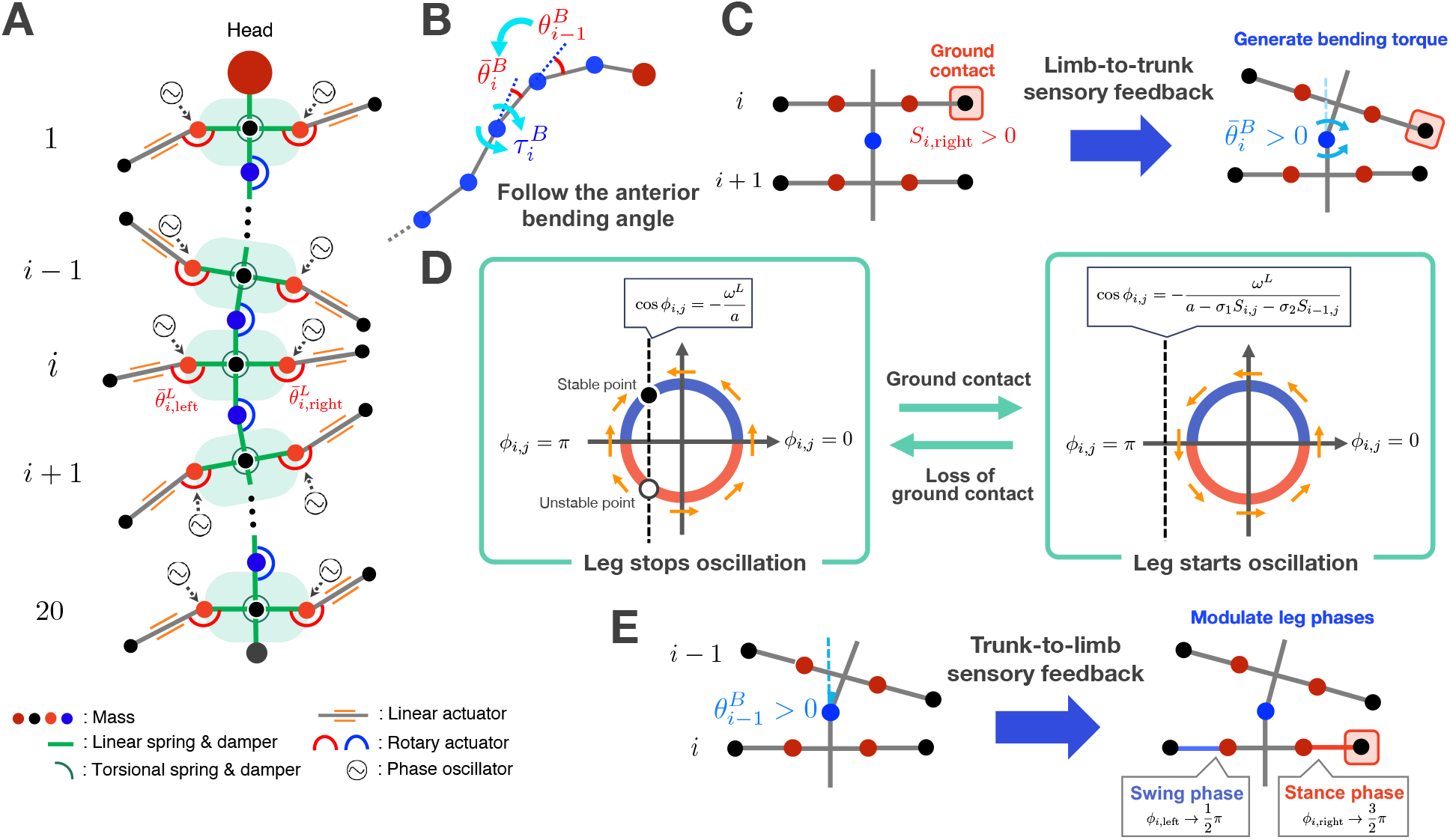
Schematic overview of the proposed model. (A) Mechanical model of the centipede body. A rotary actuator at each leg base swings the leg forward and backward, while a linear actuator adjusts the leg length solely during the stance phase. Rotary actuators between body segments enable trunk bending motion. The dots represent point masses. (B) Sensory feedback for intersegmental body coordination. Local bending torque 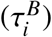 is generated so that the angle of the segment 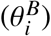 follows the angle of its nearest anterior segment 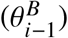, propagating the body bending posteriorly. (C) Limb-to-trunk sensory feedback for trunk–limb coordination. When a leg detects ground contact, a bending torque is generated to curve the trunk into a concave shape toward the grounded leg. (D) Sensory feedback for inter-limb coordination. When a leg and its nearest anterior legs lose contact with the ground, the leg tends to stay in a stable position and stops oscillating (left panel). Upon ground contact of either leg, the increased sensory input causes the stable and unstable solutions to disappear, and the leg starts periodic motion (right panel). (E) Trunk-to-limb sensory feedback for trunk–limb coordination. Local trunk bending modulates the phase of the leg on the concave side to be in the stance phase and that on the convex side to be in the swing phase.

Upon this lower locomotor circuit for trunk motion, we newly propose and implement the hierarchical control structure of higher centers (**‘*double inhibition*’ hypothesis**) as follows:

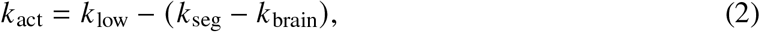

where *k*_low_ denotes the neural activation level for the trunk oscillation which can be induced by the lower nervous system, and *k*_seg_ and *k*_brain_ denote the level of inhibitory signals from the SEG and brain, respectively. *k*_act_ corresponds to the gain value of trunk bending control (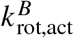 in Eq. 14) and can be varied according to the descending modulation from the higher centers. Thus, *k*_act_ is expected to be a crucial control parameter that can affect the resulting wavelength and amplitude of the trunk undulation.

#### Control circuits for leg motion

To implement the hierarchical interactions between the descending signals from the brain and SEG into the leg control circuits (**‘*leg folding*’ hypothesis**, Fig. 3), we introduced variable *H*_*i, j*_ (*i* = 1, …, *N*) which denotes the locomotor mode of the *i*th leg at the *j* side (left or right). *H*_0_ represents the descending signal from the higher centers for leg folding and is determined as follows:

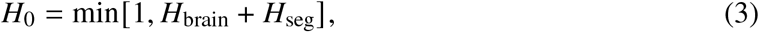

where *H*_brain_ and *H*_seg_ denote the signals from the brain and SEG, respectively. These signals are binary and represent the presence (1) or absence (0) of the descending command for leg folding. The variable *H*_*i, j*_ ranges from 0 to 1, where *H*_*i, j*_ = 0 represents the walking mode (i.e., the leg is unfolded) and *H*_*i, j*_ = 1 represents the swimming mode (i.e., the leg is folded along the side of the trunk). Note that when *H*_*i, j*_ takes an intermediate value (e.g., *H*_*i, j*_ = 0.5), the neutral position of the leg shifts backward in comparison with the walking mode, which corresponds to a transient state between walking and swimming. We described the time evolution of *H*_*i, j*_ with the following equations:

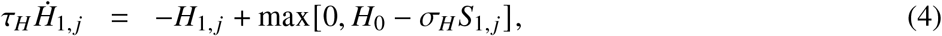

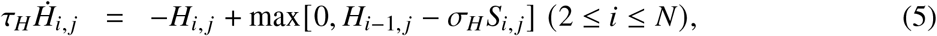

where *τ*_*H*_ is the time constant and *S*_*i, j*_ (0 ≤ *S*_*i, j*_ ≤ 1) denotes the activity level of the mechano-sensory neurons that burst when the *i*th leg in the *j* side detects a force larger than the threshold. Our model assumes that the legs, when in water, are subject to below-threshold forces (*S*_*i, j*_ = 0) and *S*_*i, j*_ becomes positive only when they receive a reaction force from the ground (see Eq. 15). Thus, Eq. 5 means that the locomotor mode of the *i*th leg converges to swimming mode (*H*_*i, j*_ → 1) only when the legs receive a descending signal for leg folding from its anterior segment (*H*_*i*−1_ = 1) and when there is no sensory input from the leg (*S*_*i, j*_ = 0).

Using the variable *H*_*i, j*_, the target joint angle of each leg 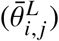 is controlled as follows:

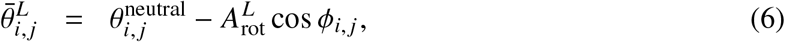

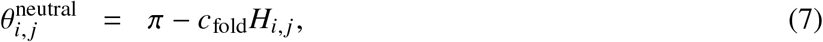

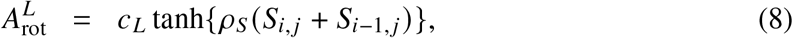

where 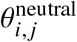 is the neutral angle of the leg, 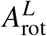 is the amplitude of leg swinging motion, *ϕ*_*i, j*_ is the neural oscillator phase of the leg, and *c*_fold_, *c*^*L*^, *ρ*_*L*_ are the positive constants. Equation 7 means that the leg unfolds during walking mode (*H*_*i, j*_ ≈ 0) and folds along the side of the body during swimming mode (*H*_*i, j*_ = 1). Additionally, Eq. 8 represents that the leg amplitude becomes positive only when the leg and/or its nearest anterior leg detect ground contact.

For the flexible inter-limb coordination during walking, here we implement the leg controllers such that the phase relationship between the legs can be formed through the local sensory feedback (**‘*ground contact*’ hypothesis**). The time evolution of oscillator phases at the *i*th leg in the left (*ϕ*_*i*,left_) and right side (*ϕ*_*i*,right_) was described as follows:

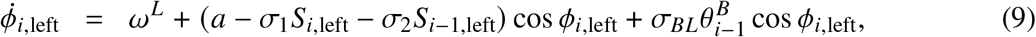

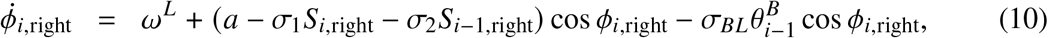

where *ω*_*L*_ is the intrinsic angular frequency and a (> *ω*^*L*^), *σ*_1_, *σ*_2_ and *σ*_*BL*_ are positive constants, *S*_*i*,left_ and *S*_*i*,right_ are mechano-sensory inputs detected at the *i*th legs (Eq. 15), and 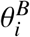 is the bending angle in the yaw direction between the (*i* − 1)th and *i*th body segment.

The second term in the right-hand side of Eqs. 9 and 10 is a sensory feedback rule proposed in the previous study (*34*) and can self-organize the traveling wave of leg movement in centipedes. Originally, the effect of sensory feedback was designed based on the behavioral finding where a centipede walking over a gap in the substrate exhibits a stop of the periodic walking motion of legs positioned over the gap until their anterior legs contact with the ground on the other side of the gap (*33*). The physical meaning of this term can be explained as follows (Fig. 4D): Suppose, for simplicity, that the body trunk is straight (*θ*_*i*_ = 0). When the leg (*i*th leg) and its nearest anterior leg ((*i* − 1)th leg) detect no ground contact (*S*_*i, j*_ = *S*_*i*−1, *j*_ = 0), 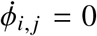 has two solutions, where one is stable and the other is unstable. Thus, the leg tends to stay at the position corresponding to the stable solution. In contrast, when the leg or its nearest anterior leg makes contact with the ground and the mechano-sensory inputs (*S*_*i, j*_, *S*_*i*−1, *j*_) have increased, it becomes *ω*^*L*^ > a − *σ*_1_*S*_*i, j*_ − *σ*_2_*S*_*i*−1, *j*_, and the stable and unstable solutions disappear. Thus, the leg starts periodic movement.

The third term on the right-hand side of Eqs. 9 and 10 is a sensory feedback from the trunk to the limb, as proposed in the previous study (*36*). As shown in Fig. 4E, the term works such that the oscillator phase of the leg at the concave side of the trunk converges to 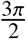, which means that the leg tends to be in the stance phase. We expect that this local sensory feedback, based on the proprioceptive information of the trunk, enables the legs to establish appropriate coordination with the trunk (‘***trunk–limb*’ hypothesis**), thereby leading to a longer stride.

### Simulation experiments

#### Reproduction of the locomotor repertoire under various neural lesions

To validate our proposed model, we first tested whether it can reproduce the behaviors of brainless and headless centipedes on land and in water as observed in our behavioral experiments. Specifically, we only changed the descending signals from the higher centers, related to the magnitude of trunk bending (i.e., *k*_brain_, *k*_seg_) and leg folding for swimming (*H*_brain_, *H*_seg_), and used common parameters for decentralized control in the lower locotomor circuits in all simulation experiments. Figure 5 shows snapshots and spatio-temporal plots of trunk bending and leg swinging motions of the simulated centipedes.

**Figure 5:**
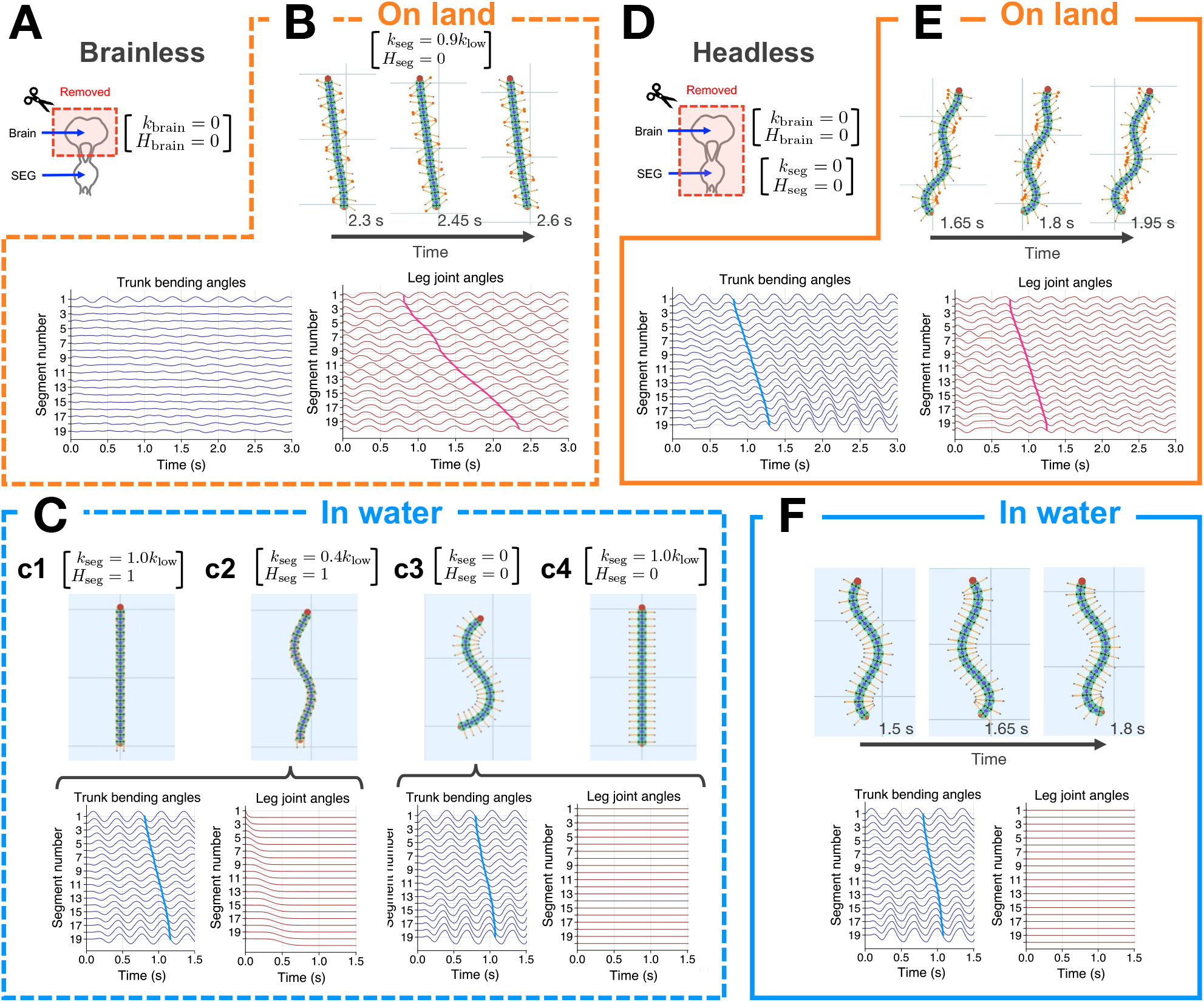
Simulated brainless and headless centipedes using the proposed model. (A and D) Schematics of the disconnected head ganglia. Descending control parameters for trunk undulation (*k*_brain_, *k*_seg_) and leg folding (*H*_brain_, *H*_seg_) are indicated across the panels. (B, C, E, and F) Snapshots and spatio-temporal plots of trunk undulation and left leg swinging for the brainless (B, C) and headless (E, F) models on land (B, E) and in water (C, F). In the simulation snapshots, leg tips in red represent the stance phase.

First, we simulated the brainless centipede by removing brain signals to the SEG (i.e., *k*_brain_ = 0, *H*_brain_ = 0; Fig. 5A, Supplementary Movie S3). On land, we assumed that SEG is active in controlling trunk motion, sending the inhibitory signal (*k*_seg_ *≃ k*_low_), while not active for leg folding (*H*_seg_ = 0). The simulated brainless centipede exhibited walking with a straight body posture and traveling wave pattern of leg movement (Fig. 5B). As we mentioned in the section on behavioral experiment, we supposed that the behavioral variations of brainless centipedes in water may be induced by the change in activity level of the SEG due to noisy inputs given by the invasive surgery. Based on this idea, in water, we assumed that SEG is normally active for inhibiting trunk motion (*k*_seg_ *≃ k*_low_) and initiating leg folding (*H*_seg_ = 1), while these two states can vary. Using four sets of SEG states, our model could reproduce all combinations of trunk and leg motions observed in our behavioral experiment (Fig. 5C): 1) suppressed trunk bending with leg folding when *k*_seg_ *≃ k*_low_, *H*_seg_ = 1 (Fig. 5c1), 2) rhythmic trunk bending with leg folding when *k*_seg_ *≃* 0, *H*_seg_ = 1 (Fig. 5c2), 3) rhythmic trunk bending with legs unfolded when *k*_seg_ *≃* 0, *H*_seg_ = 0 (Fig. 5c3), 4) suppressed trunk bending with legs unfolded when *k*_seg_ *≃ k*_low_, *H*_seg_ = 0 (Fig. 5c4).

Next, we simulated the headless centipede by removing all brain and SEG signals (i.e., *k*_brain_ = *k*_seg_ = *H*_brain_ = *H*_seg_ = 0; Fig. 5D, Supplementary Movie S4). On land, the simulated headless centipede exhibited walking with a traveling wave of leg movement and trunk bending (Fig. 5E). Furthermore, the legs were in the stance phase on the concave side of the trunk undulation, and the wave number of the leg and trunk movmevent decreased compared to that of brainless walking. In water, the simulated headless centipede exhibited a traveling wave of trunk bending with large amplitude, while the legs maintained the extended and paused position (Fig. 5F). Thus, these coordination patterns between the legs and trunk were qualitatively the same as observed in the centipedes.

In addition, we tested the capacity of our model to explain the behaviors of centipedes whose ventral nerve cord was transected in the middle of the body, which was reported in the previous study (*25*). Here, we cut the connection of neural control between the 10th and 11th segments in the model (Fig. 6A, Supplementary Movie S5). More specifically, we asssumed that the 10th trunk joint was completely passive (i.e., no active bending) and the 11th left and right legs could not recieve any descending signals (*k*_brain_ = *k*_seg_ = *H*_10, *j*_ = 0) and sensory inputs and feedback from the anterior segments(*S*_10, *j*_ = 0, *σ*_*BL*_ = 0). Descending signals from the higher centers were set as: *k*_brain_ = 0.4*k*_low_ and *k*_seg_ = *k*_low_, where *H*_brain_ = *H*_seg_ = 0 for terrestrial and *H*_brain_ = *H*_seg_ = 1 for aquatic environments. On land, the simulated centipede exhibited coordinated walking patterns in both anterior and posterior body sections (Fig. 6B). In water, the anterior section of the simulated centipede exhibited swimming with trunk undulation and leg folding, whereas the posterior section showed little trunk movement, with the legs being paused in the unfolded position (Fig. 6C). These behaviors qualitatively match those of nerve-cord-transected centipedes (*25*).

**Figure 6:**
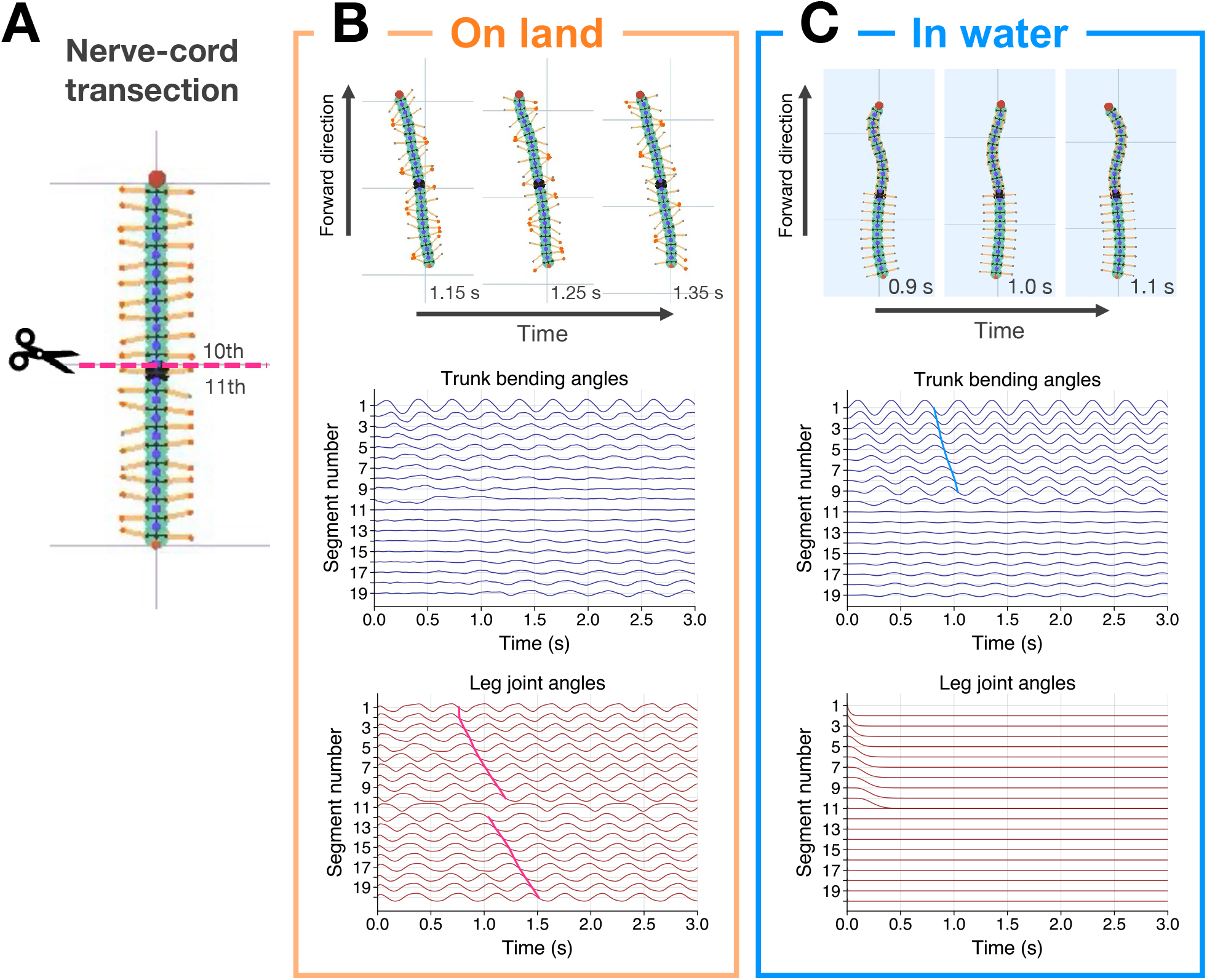
Simulated nerve-cord-transected centipedes using the proposed model. (A) Schematic of the nerve-cord transection. The connection of neural control between the 10th and 11th segments was cut. (B and C) Snapshots and spatio-temporal plots of trunk undulation and left leg swinging on land (B) and in water (C). In the simulation snapshots, red leg tips represent the stance phase, and black segments indicate the segments immediately posterior to the nerve-cord transection site.

#### Speed-dependent gait transition on land

We examined whether our model could reproduce the gait transitions between slow and fast walking in centipedes (Supplementary Movie S6). Although various factors may induce such transitions, we tested whether the change in the magnitude of the descending brain signal for trunk undulation (*k*_brain_) is sufficient to elicit them. For simplicity, we assumed that the SEG is kept active for trunk undulation, sending a constant inhibitory signal (*k*_seg_ = *k*_low_), and varied *k*_brain_ in a stepwise manner (low to high, and back to low) during simulated terrestrial locomotion. Descending signals for leg folding (*H*_brain_, *H*_seg_) were set to be zero.

Representative snapshots of the simulated centipede were shown in Fig. 7A. Figs. 7B and C present the spatio-temporal plots of trunk bending and leg swinging motions, respectively, accompanied by their corresponding descending control signals. The simulated centipede initially exhibited a slow walking gait with an almost straight trunk posture under low *k*_brain_. Upon the increase of *k*_brain_ at 3.0 s, the centipede transitioned to a fast walking gait characterized by large trunk undulations, where the legs were in the stance phase on the concave side of the trunk. Subsequently, when *k*_brain_ was reset to the low level at 6.0 s, the gait returned to slow walking pattern. Quantitative evaluation across 20 trials with random initial leg oscillator phases confirmed that: (i) the higher value of *k*_brain_ resulted in increased locomotion speeds (Fig. 7D), and (ii) the proposed control circuit autonomously self-organized distinct gaits, ranging from slow walking with large inter-limb phase differences (i.e., large wave numbers) to fast walking with small phase differences (i.e., small wave numbers) (Fig. 7E). These results demonstrate that our model successfully reproduces adaptive trunk–limb and inter-limb coordination in response to the magnitude of trunk undulation, associated with changes in locomotion speed as observed in real centipedes (*26*).

**Figure 7:**
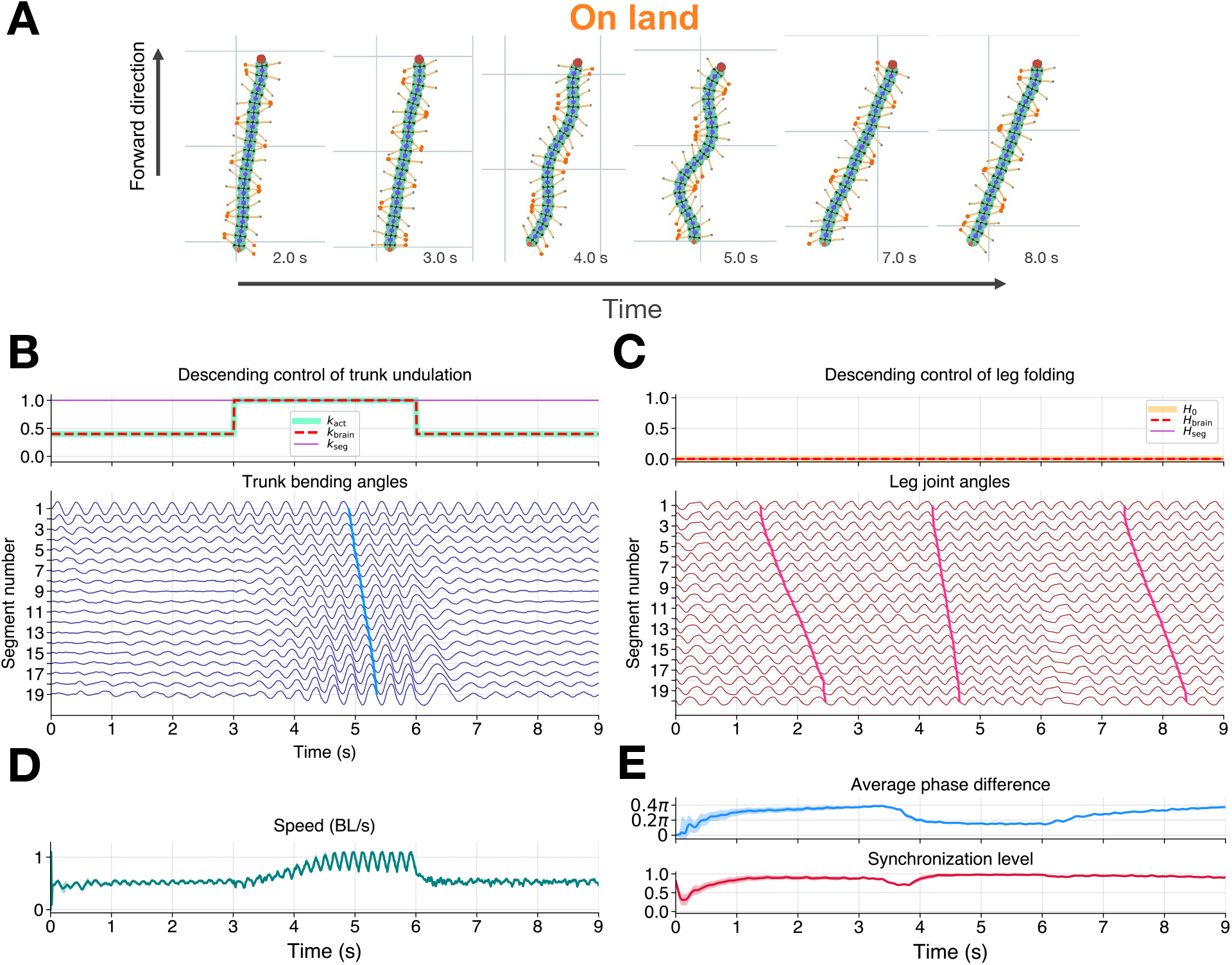
Spontaneous terrestrial gait transitions of the simulated centipede using the proposed model. (A) Snapshots of the simulated centipede on land. Leg tips in red represent the stance phase. (B) Time evolution of descending control parameters for trunk undulation (*k*_brain_, *k*_seg_, *k*_act_) normalized by *k*_low_, and spatio-temporal plots of trunk undulation. (C) Time evolution of descending control parameters for leg folding (*H*_brain_, *H*_seg_, *H*_0_) and spatio-temporal plots of left leg swinging. (D) Center-of-mass speed normalized by body length (BL). (E) Time evolution of the average phase difference between successive legs on the left side and the synchronization level of the leg phase differences (see Materials and Methods). Solid lines and shaded areas in (D and E) indicate the mean and standard deviation for 20 trials starting from random initial leg oscillator phases. A synchronization level of 1.0 indicates that the oscillator phases of successive legs are perfectly synchronized with a constant phase difference, while values approaching 0.0 indicate randomly distributed phase differences.

#### Walk-swim transition in response to environmental changes

We examined whether our model could also reproduce the flexible transition between walking and swimming in response to environmental changes (Supplementary Movie S7), as reported in real centipedes (*25*). In the simulations, we prepared an arena with a land–water–land configuration. We assumed that when the head of the simulated centipede entered the water, the descending signals for both trunk undulation (*k*_brain_) and leg folding (*H*_brain_, *H*_seg_) were switched to higher values to initiate swimming behavior. Subsequently, upon the head’s re-entry onto land, only the leg folding signals (*H*_brain_, *H*_seg_) were deactivated, while *k*_brain_ was maintained at the high level, allowing the centipede to transition to a fast walking gait.

Representative snapshots of the simulated centipede were shown in Fig. 8A. Figs. 8B and C present the spatio-temporal plots of trunk bending and leg swinging motions, respectively, accompanied by their corresponding descending control signals. At the beginning of the simulation, the centipede walked on land by propagating a traveling wave of leg movement while keeping the trunk almost straight. As the anterior body section entered the water, it immediately started to shift to swimming, characterized by rhythmic trunk undulation and leg folding. In contrast, the posterior section remaining on land continued walking (see the snapshot of 2.96 s in Fig. 8A). This behavior was generated by local sensory feedback from the legs, which overrode the leg-folding signals from higher centers. Once the entire body was submerged, the simulated centipede exhibited a full swimming motion. Subsequently, when transitioning from water to land, the anterior section immediately shifted to walking, while the posterior legs remaining in the water gradually began to unfold (see the snapshot of 4.56 s in Fig. 8A). This locomotor transition was generated by the proposed control mechanism, in which each leg mimics the state of the leg preceding it (Eq. 5). Upon entering the land area, once a posterior leg detected ground contact, it initiated rhythmic motion and established a coordinated walking gait. Since the descending signal for the magnitude of trunk undulation (*k*_brain_) was maintained at a high level, the centipede exhibited a fast walking gait with large trunk undulations after reaching land. Thus, the proposed model could reproduce flexible locomotor transitions between walking and swimming similar to real centipedes (*25*).

**Figure 8:**
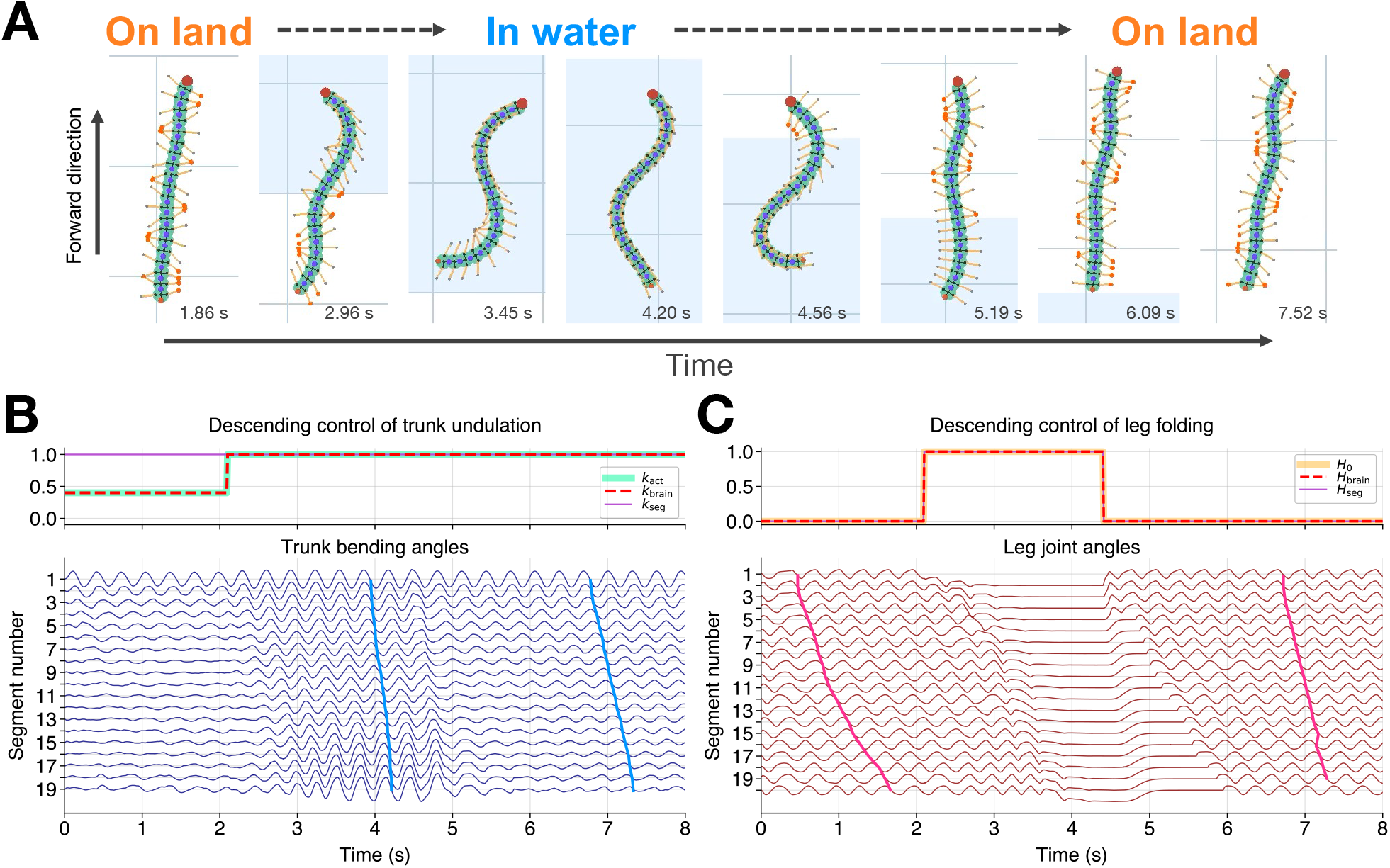
Adaptive walk-swim transition of the simulated centipede using the proposed model. (A) Snapshots of the simulated centipede crossing a land-water-land environment. White and light blue areas represent land and water, respectively. Leg tips in red represent the stance phase. (B) Time evolution of descending control parameters for trunk undulation (*k*_brain_, *k*_seg_, *k*_act_) normalized by *k*_low_, and spatio-temporal plots of trunk undulation. (C) Time evolution of descending control parameters for leg folding (*H*_brain_, *H*_seg_, *H*_0_) and spatio-temporal plots of left leg swinging.

## Discussion

Many neurobiological and modeling studies have claimed the importance of the interplay between the higher centers and lower locomotor circuits in adaptive locomotion (*15, 41–45*). However, most previous studies focused on the coordination between homogeneous body parts such as inter-limb coordination during walking in legged animals (*46–49*) and intersegmental body coordination during swimming and crawling in limbless animals (*50–55*). The key challenge is elucidating an integrative model that can account for a diverse locomotor repertoire, including qualitatively different coordination patterns between heterogenous body parts (e.g., trunk–limb coordination), and flexible transitions between them. In this study, leveraging the versatile locomotion and robustness against neural lesion of centipedes, we could estimate the contributions of descending signals from the higher centers through living animal experiments. By combining these findings with computational modeling, we obtained an integrative view of how lower locomotor circuits with descending commands, manage a wide variety of locomotor patterns in a situtation-dependent manner.

Our behavioral experiments using headless centipedes provide empirical evidence supporting the hypotheses on the walking control, proposed in our computational modeling studies. The ‘*trunk–limb*’ hypothesis (*36*) was behaviorally validated by the observation that headless centipedes exhibited a fast walking pattern on land, where trunk and limb movements were coordinated as in intact animals. This persistence suggests that the trunk–limb coordination can be managed within the lower locomotor circuits. Furthermore, the ‘ground-contact’ hypothesis (*33, 34*) was also supported by the fact that headless centipedes exhibited robust inter-limb coordination on land but failed to initiate any rhythmic leg motion in water. This behavioral contrast between terrestrial and aquatic environments indicates that the leg mechanosensory feedback from the ground is crucial for both activating neural oscillators controlling the legs and establishing inter-limb coordination.

Step-wise removal of the brain and SEG further informed us how descending control interacts with lower locomotor circuits to achieve versatile locomotion. Our results revealed that even in the absence of higher centers, the lower circuits possess the intrinsic capability to generate rhythmic motions in both the legs and the trunk. In contrast, specific coordinated behaviors observed in intact animals, such as the folding of legs during swimming or the maintenance of a straight trunk during slow walking, were rarely observed in headless centipedes. This suggests that these behaviors are actively induced by descending signals that suppress the autonomous rhythmic potential of the lower circuits (‘*leg folding*’ and ‘*double inhibition*’ hypotheses). Rather than explicitly controlling the complex coordination among the vast degrees of freedom in the body, the higher centers may achieve flexible multimodal locomotion by selectively releasing or intensifying this inhibition. Using the neuro-mechanical model of centipedes, we demonstrated that locomotor transitions between slow walking, fast walking, and swimming can be achieved only by changing a few descending signals controlling the magnitude of trunk undulation and/or leg folding (Figs. 7 and 8). Thus, the harnessing strategy we found in the descending control exploits the self-organization of the locomotor patterns driven by the lower locomotor circuits via sensory feedback, and provides a computationally simple mechanism to flexibly orchestrate diverse locomotor patterns in a situation-dependent manner.

The difference in the behavioral findings between this study and our previous one using ventral-nerve-cord-transected centipedes (*25*) also has implications on the heterogeneity of lower locomotor circuits (segmental circutis). We previously reported that when the connectives between the segmental ganglia at the middle section of the body, the posterior half of the body could not initiate the swimming (*i*.*e*., leg folding and rhythmic trunk undulation). In contrast, this study reported that the headless centipedes, athough showed similar inability to fold their legs, were still able to generate swimming-like rhythmic tunk undulation. We hypothesize that behavioral differences arise from differences in transection position and that the segmental ganglia in the posterior half of the animal may have less sensory driven trunk oscillation excitability when descending drive is not present. Based on this hypothesis (that body segment position affects segment oscillation capacity; the ‘segment heterogeneity’ hypothesis), our current model assumes intrinsic oscillation exists solely in the first anterior segment for simplicity. With this assumption, our model successfully replicates both transected swimming (no body oscillation post transection, Fig. 6) and decapitated aquatic movement (body oscillation present in water, Fig. 5). Ongoing real animal experiments that test the segment heterogeneity hypothesis will be essential to understand how body segment circuitry differs along the body and contributes to adaptive locomotor function in centipedes.

## Materials and Methods

### Behavioral experiments

All centipedes (*Scolopendra subspinipes mutilans*) were wild-caught in Wakayama, Japan, and a total of 15 individuals were used for behavioral observations. To examine the contribution of descending signals from the higher centers to locomotor behavior, we prepared two experimental groups: brainless and headless centipedes. For the brainless condition, nine centipedes were used, with a body length of 8.58 ± 0.70 cm (mean ± SD). The remaining six centipedes were used for the headless condition, with a body length of 11.78 ± 0.71 cm (mean ± SD).

The brain was removed through the following surgical procedure: each centipede was anesthetized with CO_2_ and placed on an ice-cold plate to prevent awakening. A scalpel was used to open the head cuticle, and the connectives between the brain and SEG were cut. To avoid damage to internal tissues, the cuticle was reattached to its original position. For the headless condition, the head was removed just below the first leg-bearing segment using microscissors after CO_2_ anesthesia. Each specimen was kept in a warm environment (23–25 °C) for 0.5–1.0 h to allow recovery before observations. All experiments were recorded from the top view using high-speed video cameras (DITECT, HAS-U2). Except for the experiments on land using headless centipedes (300 fps; Fig. 2E), all videos were recorded at a frame rate of 150 fps (Fig. 2B, C, F).

### Model

The linear and rotational actuators implemented at each leg and trunk generate forces and torques to achieve the target positions based on proportional-derivative control. The target leg length during swing and stance phases was described by:

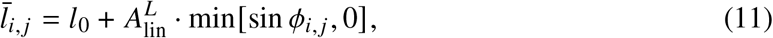

where *l*_0_ is the constant leg length during swing phase (sin *ϕ*_*i, j*_ > 0) and 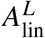 is a positive constant that defines the shortening of the leg length during stance phase (sin *ϕ*_*i, j*_ < 0). The linear actuator at the leg generates force (*f*_*i, j*_) as follows:

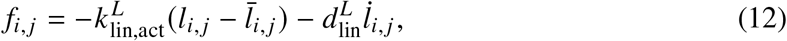

where *l*_*i, j*_ is the actual leg length, 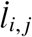 denotes its time derivative, and 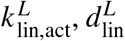 are the proportional and derivative gain values. The rotational actuators at the leg and trunk generate the torque 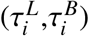 determined by the following equations:

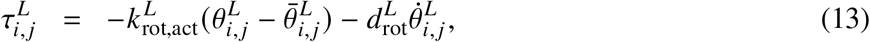

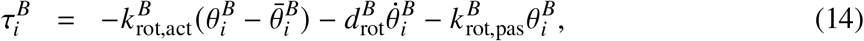

where 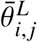 and 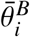 are the target angles of the leg and trunk joints, respectively, and 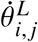 and 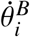 denote the time derivatives of each actual angle. 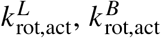 are the proportional gains, and 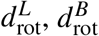 are the derivative gains. Note that the passive torsional spring is implemented at the trunk joint to maintain the body stiffness 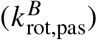 independent of the activity level of the trunk undulation controlled by the descending signal (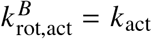, Eq. 2).

The interaction forces between the body and environment are described as viscous friction for simplicity. Each mass point receives viscous frictional force, proportional to its velocity, from the environment (the coefficients of viscous friction are summarized in Table S1). When the body is on land, we assumed that the leg tip lifts off the ground during 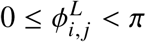 and makes contact with the ground during 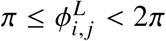. Thus, the friction coefficient of the leg tip mass during 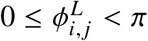 is set to be zero. In addition, the friction coefficient of the body trunk masses is set to be zero since the body trunk of a real centipede mostly lifts off the ground during walking. Meanwhile, when the body is in water, the whole body always interacts with the environment, and thus, the friction coefficients have non-zero values. We assumed that the friction coefficient of the leg tip masses in water (*μ*_*w*_) is smaller than that on the ground (*μ*_*g*_). Moreover, it is also assumed that the friction coefficient of the body trunk masses in the tangential direction (i.e. forward-backward direction), *μ*_*t*_ is set to be smaller than that in the normal direction (i.e. lateral direction), *μ*_*n*_.

We have described the activity level of mechano-sensory neurons in each leg (*S*_*i, j*_, *j* ∈ {left, right}) as follows:

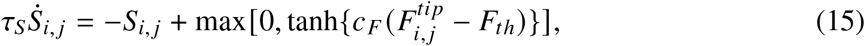

where *τ*_*s*_ is the time constant, *c*_*F*_ is the positive constant, 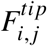 is the reaction force from the environment (i.e. viscous friction) received at the leg tip, and *F*_*th*_ is the threshold value. We assumed that inside water, the leg receives a small force under the threshold value (i.e. *S*_*i, j*_ = 0) and *S*_*i, j*_ becomes positive only when the leg receives reaction force from the ground.

### Simulation experiments

The program for simulation was written in C++ and the simulation results were visualized using OpenGL. The differential equations were solved using the fourth-order Runge-Kutta method with a time step of 1.0 × 10^−5^ s. The physical parameters of the simulated centipede were set to match those of real animals: the total body length and mass were 10.0 cm and 3.0 g, respectively. The leg length and the mass of each leg tip were set at 0.75 cm and 2.8 mg, respectively. The parameter values employed in the simulations were listed in Tables S1 and S2. For the first body segment, *σ*_1_, *σ*_3_, and *S*_*i*−1, *j*_ were set to be zero. At the initial condition of all experiments, the simulated centipede had a straight body posture and the leg-tip positions were set randomly by varying the initial phases of the leg oscillators.

The average phase difference between the neighboring leg oscillators (Δ*ϕ*_ave_, shown in Fig. 7E) can be calculated as follows:

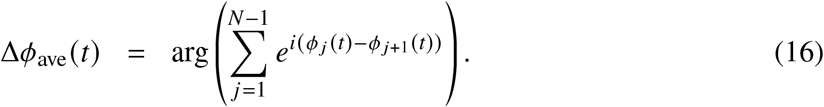

The phase difference ranges from −*π* to *π*. When Δ*ϕ*_ave_ is positive, the phase of each leg is advancing compared to that of its nearest posterior leg on average, which means that the traveling wave of the leg motion propagates from anterior to posterior. In addition, the synchronization level of the leg oscillations (shown in Fig. 7E) can be measured by the following metric:

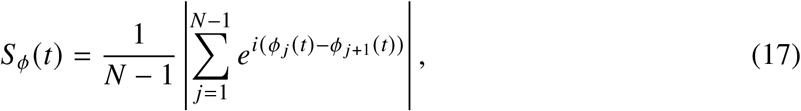

where *N* is the leg number. The synchronization level *S*_*ϕ*_ (*t*) of 1.0 indicates that the oscillator phases of adjacent legs are completely synchronized with a constant phase difference, while the value approaches 0.0 when the phase differences are randomly distributed.

## Funding

E.M.S. and A.I. were funded by Human Frontier Science Program (grant RGP0027/2017). K.Y., H.A., and A.I. were funded by the JSPS KAKENHI (JP22H00216). K.Y., T.K., and A.I. were funded by the JSPS KAKENHI (JP23KK0072). K.Y. was funded by the JSPS KAKENHI (JP25K03201).

## Author contributions

Conceptualization: K.Y., E.M.S., A.I., Methodology: K.Y., E.M.S., T.K., H.A., Software: K.Y., Validation: K.Y., Formal analysis: K.Y., Investigation: K.Y., E.M.S., H.A., Resources: K.Y., H.A., A.I., Data curation: K.Y., Writing – original draft: K.Y., E.M.S., Writing – review & editing: K.Y., E.M.S., T.K., H.A., A.I., Visualization: K.Y., Supervision: E.M.S., T.K., A.I., Funding acquisition: K.Y., E.M.S., T.K., H.A., A.I.

## Competing interests

There are no competing interests to declare.

### Data and materials availability

All data needed to evaluate the conclusions in the paper are present in the paper and/or the Supplementary Materials.

## Supplementary materials

**Table S1:** Body and environment parameters used in the simulations.

| Parameter | Value | Dimension |
| --- | --- | --- |
| (Body parameters) |  |  |
| Number of body segments | 20 | - |
| Body segment length | $0.5 \times 10^{-2}$ | [m] |
| Body segment width | $0.5 \times 10^{-2}$ | [m] |
| Body segment mass | $1.4 \times 10^{-4}$ | [kg] |
| Leg length ($l_0$) | $0.75 \times 10^{-2}$ | [m] |
| Leg tip mass | $2.8 \times 10^{-6}$ | [kg] |
| Total body length | 0.1 | [m] |
| Total body mass | $3.0 \times 10^{-3}$ | [kg] |
| $k_{\text{lin,act}}^L$ | 2.8 | $[\text{kg} \cdot \text{s}^{-2}]$ |
| $d_{\text{lin}}^L$ | $2.8 \times 10^{-2}$ | $[\text{kg} \cdot \text{s}^{-1}]$ |
| $k_{\text{rot,act}}^L$ | $4.87 \times 10^{-4}$ | $[\text{kg} \cdot \text{m}^2 \cdot \text{s}^{-2}]$ |
| $d_{\text{rot}}^L$ | $4.87 \times 10^{-7}$ | $[\text{kg} \cdot \text{m}^2 \cdot \text{s}^{-1}]$ |
| $k_{\text{rot,act}}^B$ | $4.87 \times 10^{-3}$ | $[\text{kg} \cdot \text{m}^2 \cdot \text{s}^{-2}]$ |
| $k_{\text{rot,pas}}^B$ | $4.87 \times 10^{-5}$ | $[\text{kg} \cdot \text{m}^2 \cdot \text{s}^{-2}]$ |
| $d_{\text{rot}}^B$ | $2.43 \times 10^{-6}$ | $[\text{kg} \cdot \text{m}^2 \cdot \text{s}^{-1}]$ |
| (Environment parameters) |  |  |
| $\mu_g$ | $1.4 \times 10^{-2}$ | $[\text{kg} \cdot \text{s}^{-1}]$ |
| $\mu_w$ | $2.8 \times 10^{-5}$ | $[\text{kg} \cdot \text{s}^{-1}]$ |
| $\mu_t$ | $2.8 \times 10^{-4}$ | $[\text{kg} \cdot \text{s}^{-1}]$ |
| $\mu_n$ | $2.8 \times 10^{-3}$ | $[\text{kg} \cdot \text{s}^{-1}]$ |

**Table S2:** Neural control parameters used in the simulations.

| Parameter | Value | Dimension |
| --- | --- | --- |
| (Trunk control parameters) |  |  |
| $\omega^B$ | 20 | $[\text{s}^{-1}]$ |
| $A^B$ | $20\pi/180$ | - |
| $\sigma_{LB}$ | 0.1 | - |
| (Leg control parameters) |  |  |
| $\omega^L$ | 20 | $[\text{s}^{-1}]$ |
| $a$ | 22 | $[\text{s}^{-1}]$ |
| $\sigma_1$ | 15 | $[\text{s}^{-1}]$ |
| $\sigma_2$ | 30 | $[\text{s}^{-1}]$ |
| $\sigma_{BL}$ | 60 | $[\text{s}^{-1}]$ |
| $c_L$ | $35\pi/180$ | - |
| $\rho_S$ | 10 | - |
| $A_{\text{lin}}^L$ | $1.25 \times 10^{-3}$ | $[\text{m}]$ |
| (Sensory information parameters) |  |  |
| $\tau_S$ | 0.157 | $[\text{s}]$ |
| $c_F$ | $1.0 \times 10^4$ | - |
| $F_{th}$ | $1.0 \times 10^1$ | - |
| (Descending control parameters) |  |  |
| $\tau_H$ | 0.025 | $[\text{s}]$ |
| $\sigma_H$ | 1 | - |
| $c_{\text{fold}}$ | $\pi/2$ | - |

**Caption for Movie S1. Behaviors of brainless centipedes on land and in water**.

**Caption for Movie S2. Behaviors of headless centipedes on land and in water**.

**Caption for Movie S3. Simulated brainless centipedes on land and in water**.

**Caption for Movie S4. Simulated headless centipedes on land and in water**.

**Caption for Movie S5. Simulated nerve-cord-transected centipedes on land and in water**.

**Caption for Movie S6. Spontaneous terrestrial gait transitions of the simulated centipede**.

**Caption for Movie S7. Adaptive walk-swim transition of the simulated centipede**.

## Notes

### Competing Interest Statement

The authors have declared no competing interest.

